# PROPEL: a scalable model for postbaccalaureate training to promote diversity in the biomedical workforce

**DOI:** 10.1101/2024.05.01.592104

**Authors:** Jessica Allen, Ekland Abdiwahab, Meghan D. Morris, Claude Jourdan Le Saux, Paola Betancur, K. Mark Ansel, Ryan D. Hernandez, Todd G. Nystul

## Abstract

Promoting diversity in the scientific workforce is crucial for harnessing the potential of available talent and ensuring equitable access to Science, Technology, Engineering, Mathematics, and Medicine (STEM-M) careers. We have developed an innovative program called Post-baccalaureate Research Opportunity to Promote Equity in Learning (PROPEL) that provides scientific and career development training for postbaccalaureate scholars from historically excluded backgrounds in STEM-M fields with an interest in pursuing a PhD or MD/PhD degree. Our program is distinct from other postbaccalaureate programs in that scholars are hired by individual labs rather than funded centrally by the program. This funding mechanism removes the idea that central funding is necessary to encourage faculty to train diverse scholars and allows the program to scale dynamically according to the needs of the scientific community. The PROPEL program started in 2020 with six scholars and has since grown to an enrollment of over 100, making it the largest postbaccalaureate program for biomedical research in the country. Here, we describe the program structure and curriculum, our strategy for recruitment, the enrollment trends; the program demographics; metrics of scholar engagement; and outcomes for scholars who completed the program in 2023. Our experience demonstrates the strong demand from both scholars and faculty for programming of this type and describes the feasibility of implementation.

## PERSPECTIVE

Despite an upward trend in the diversity of the Science, Technology, Engineering, Mathematics, and Medicine (STEM-M) workforce, we have yet to reach representation that matches the US population. For example, whereas Black, African American, Hispanic, Latine, American Indian, and Alaskan Native groups collectively constituted about 31% of the US population and a similar percentage of the total labor force in 2021, an NSF study estimated that these groups constituted just under 25% of the US STEM-M workforce in that year (an increase from 18% in 2011) (1). Similarly, an estimated 3% of the US STEM-M workforce in 2021 was comprised of individuals with a disability compared to 9% of the overall population. We see these disparities at the local level as well. For example, at the University of California, San Francisco (our institution), while the percentage of graduate students from underrepresented racial and ethnic groups has risen from 22% to 25% over the past five years,^1^ it does not reflect the demographics of California, where over 50% of adults in the prime working age range identify as Latine, Black, or Native American.^2^

Increasing representation would provide a wide range of benefits for STEM-M fields, especially in combination with cultural competence in the workplace and representative leadership (2). Better utilization of the entire labor pool would increase labor market efficiency and ensure more equitable access to this important sector of the economy. In addition, many studies have found that more diverse teams have increased creativity and problem-solving capability (3, 4), productivity (5), and resilience (6). Moreover, an increase in STEM-M workforce diversity is also important for keeping pace with the changing demographics of our society (7), which is particularly salient at a time when the US STEM-M fields are facing a severe labor shortage (8). Therefore, programs that promote the recruitment and retention of scholars from diverse backgrounds into STEM-M career paths at each stage of the educational pipeline are urgently needed.

PhD training is typically a highly formative period during which scholars develop the skills and expertise required to make independent contributions to their chosen field. The resulting degree is also a required credential for many of the senior positions in STEM-M career paths. However, entry into biomedical doctoral programs is very competitive, with many programs prioritizing applicants who have one or more years of full-time research experience before applying (9). While this emphasis on prior research experience helps to identify applicants who may have the experience and knowledge to succeed in a rigorous PhD program, it also creates yet another barrier that exacerbates the exclusion of some student groups from PhD programs as these students may be more likely to attend colleges that have relatively fewer resources (10, 11), and undergraduate research positions are less available. Further, while some undergraduate research positions can be paid through work-study or other funding mechanisms, many undergraduate research opportunities are unpaid (12). In addition, students from historically excluded groups may have fewer opportunities to network and integrate into academic communities during their undergraduate careers (13). Thus, many students from historically excluded groups graduate from college without the same level of skills, professional network, or credentials as students from groups with higher social capital, leading to continued disparities in STEM-M graduate program application and attendance.

To address this gap, several federally funded postbaccalaureate programs have been developed that provide 1-2 years of scientific and career development training after college for scholars from historically excluded groups. Two of the most well-established of these programs are the NIH Postbaccalaureate Intramural Training Award (IRTA) and the NIGMS PREP program, and more recent programs include PREP programs funded by other NIH institutes and the NSF Research and Mentoring for Postbaccalaureates (RAMP). Unlike Master’s programs, these postbaccalaureate programs typically provide scholars with a full-time salary and benefits, do not charge tuition, and do not award a degree. Instead, these postbaccalaureate programs are, foremost, opportunities to conduct original research in a lab or other research setting at the university. In addition, they provide scholars with scientific and professional development activities, like application workshops, information panels, community-building activities, and coursework, thus providing a more structured path toward acceptance into a graduate program.

Retrospective analyses have demonstrated that NIGMS PREP programs are highly successful (14, 15). However, because scholars in these programs are funded by training grants awarded to the university, they are only available at schools that have applied for and received one of these grants, and the size of the program is limited to the number of slots awarded on the grant. This caps the number of slots that are available for these positions nationwide, and demand for a slot in these programs far exceeds supply. One common rationale for the traditional approach of centrally funded training programs that promote diversity is that training grant slots are important to incentivize the involvement of research faculty. However, an alternative framework is an asset-based model, which recognizes and values the inherent strengths of scholars from historically excluded backgrounds (16, 17). This model asserts that scholars from these historically excluded groups would be competitive for postbaccalaureate training opportunities if given sufficient exposure to research labs looking to hire.

In 2020, we launched a postbaccalaureate research program at the University of California, San Francisco (UCSF), which we named Postbaccalaureate Research Opportunity to Promote Equity in Learning (PROPEL). Unlike traditional programs, PROPEL takes advantage of research funding from individual labs and institutional resources to support scholars. While the university provides funding for program administration and curriculum, scholars’ salaries and benefits are covered by their host laboratories. This model combines the advantages of both training grant-funded programs and independent technician hires.

Scholars engage in structured group-based professional development alongside full-time applied research. In this way, PROPEL provides similar research, professional, and scientific development opportunities as the training-grant funded programs do, but PROPEL is not involved in the initial selection process, as they must be hired into an individual research group at UCSF before joining PROPEL. This removes the enrollment cap imposed by training grant funds and allows the program to scale dynamically to fit the needs of the research community at the university. Here, we describe the program structure and curriculum; our strategy for recruitment and the enrollment trends; the program demographics; metrics of scholar engagement; and outcomes for scholars who completed the program in the past two years.

## DESIGN OF THE PROGRAM

### Program administration

The administrative structure of the PROPEL program consists of two faculty Co-Directors, a Program Administrator, and an Advisory Council that consists of the Co-Directors, the Program Administrator, and four additional faculty members who are closely involved with different aspects of the program, such as curriculum or outreach. In addition, a PROPEL Scholars Council, comprised of 4-8 scholars who are nominated and chosen by the scholars, organizes scholar-initiated activities and serves as a liaison between the scholars and the program administration.

### Enrollment

The PROPEL program is distinct from other postbaccalaureate programs in that it is designed for scholars who are working as employees in a research group. Scholars do not pay tuition, and no degree or certificate is awarded. In the first three years, the PROPEL program accepted all applicants who (1) had a Bachelor’s degree; (2) were employed full-time at UCSF; (3) would meet the suggested eligibility criteria for an NIH Research Supplement to Promote Diversity in Health-Related Research (“NIH Diversity Supplement”),^3^ whether or not they are funded by one; (4) committed to participating in the program for at least a year; and (5) expressed a strong interest in pursuing a PhD or MD/PhD degree. With these criteria, there was a 90.6% acceptance rate from the inception of the program in August, 2020 to August, 2023. Scholars were admitted into the program on a rolling basis, at any time of the year, after their applications were reviewed by at least two faculty on the PROPEL administrative team to confirm that they met the eligibility requirements. There is no limit to the length of time that a scholar can be in the program but, as described below, the curriculum is designed to be completed in two years, but all curriculum could be completed in 1 year, if necessary (**Figure 1**). Indeed, from August, 2020 to August, 2023, the mean length of time in the program among the scholars who exited was 1.73 years. Following recent changes in the enrollment policy, scholars and their UCSF faculty mentor are now required to apply during one of four admissions cycles each year and admission is based on a selection process rather than eligibility criteria alone.

**Fig. 1.**
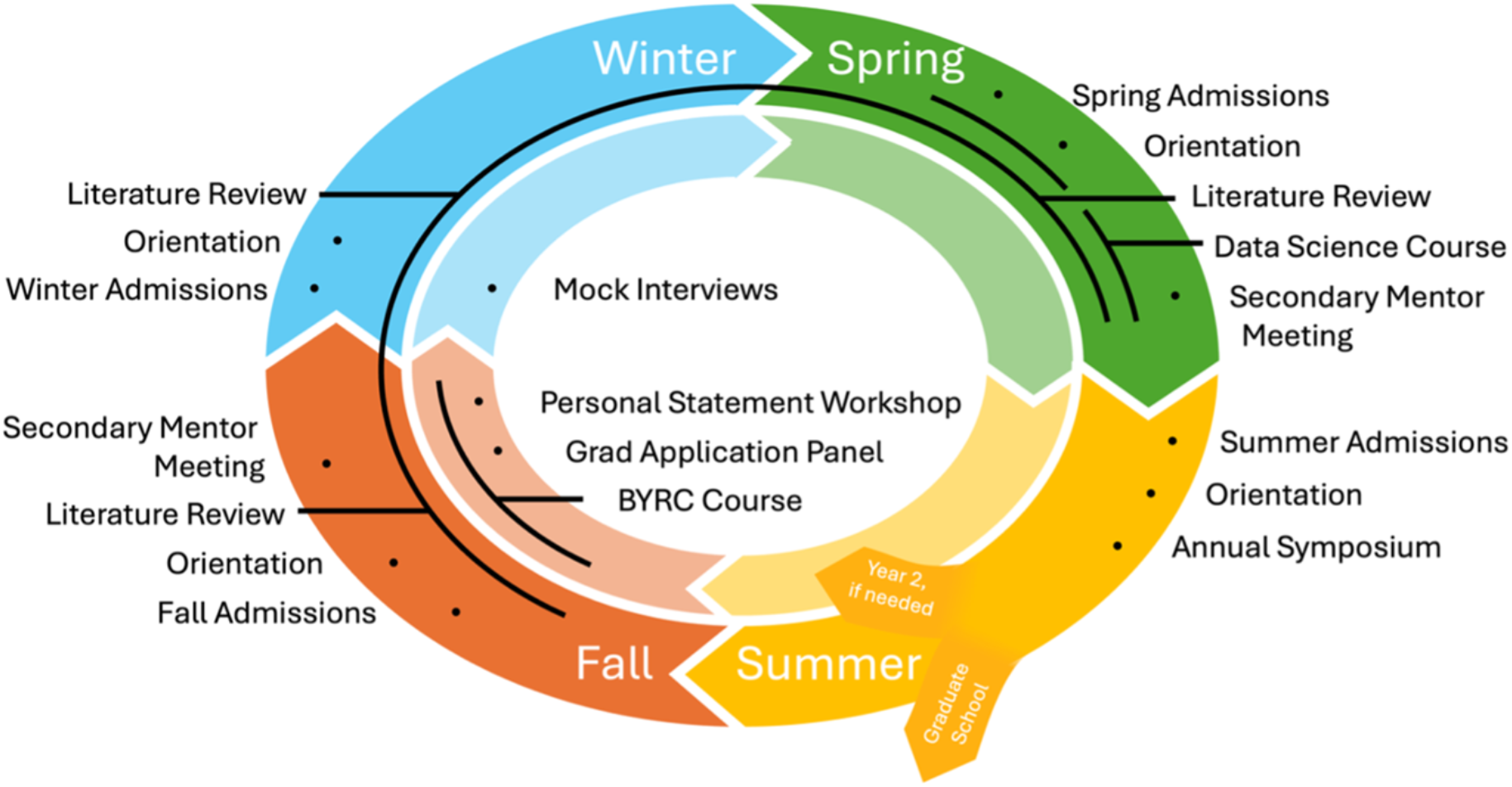
Diagram showing the cycle of activities such as admissions, courses, and workshops in the PROPEL program. The outer circle includes curriculum, while the inner circle consists of workshops and interventions designed for scholars applying to graduate school. All courses and workshops can be completed in 1 year, depending on the scholar’s application timeline.

A major route of entry into the PROPEL program has been through the UCSF NIH Diversity Supplement Matchmaking Event. This event, held annually in January or February, aims to match scholars at the postbaccalaureate level who would qualify for an NIH Diversity Supplement with UCSF faculty who are looking to hire research technicians. Before the event, both faculty and scholars fill out a questionnaire about their research interests and the scholars also submit standard application materials such as a curriculum vitae and contact information for references. After the eligibility of both the faculty and scholar applicants has been confirmed, the participant lists are distributed, and both faculty and scholars indicate whom they would like to meet at the event. This information, along with the questionnaire data about research interests, is used to match faculty and scholar candidates for four to eight 15-minute “speed interviews” on a single day. Scholars and faculty are encouraged to follow up with individuals they would be interested in working with to set up additional interviews, hopefully resulting in a new hire. Scholars are recruited to the event from around the country and interviews are conducted by video conference so that no travel is necessary to attend. Past events have attracted approximately 90 scholars and a similar number of UCSF faculty, resulting in several dozen new hires per year. As of January 2023, 51% (81/160) of the scholars who joined the PROPEL program attended a Matchmaking Event, and most of these scholars joined the lab of a faculty member they met at the event. The rapid growth in the cumulative number of scholars who have joined PROPEL since the inception of the program is shown in Figure 2.

**Fig. 2.**
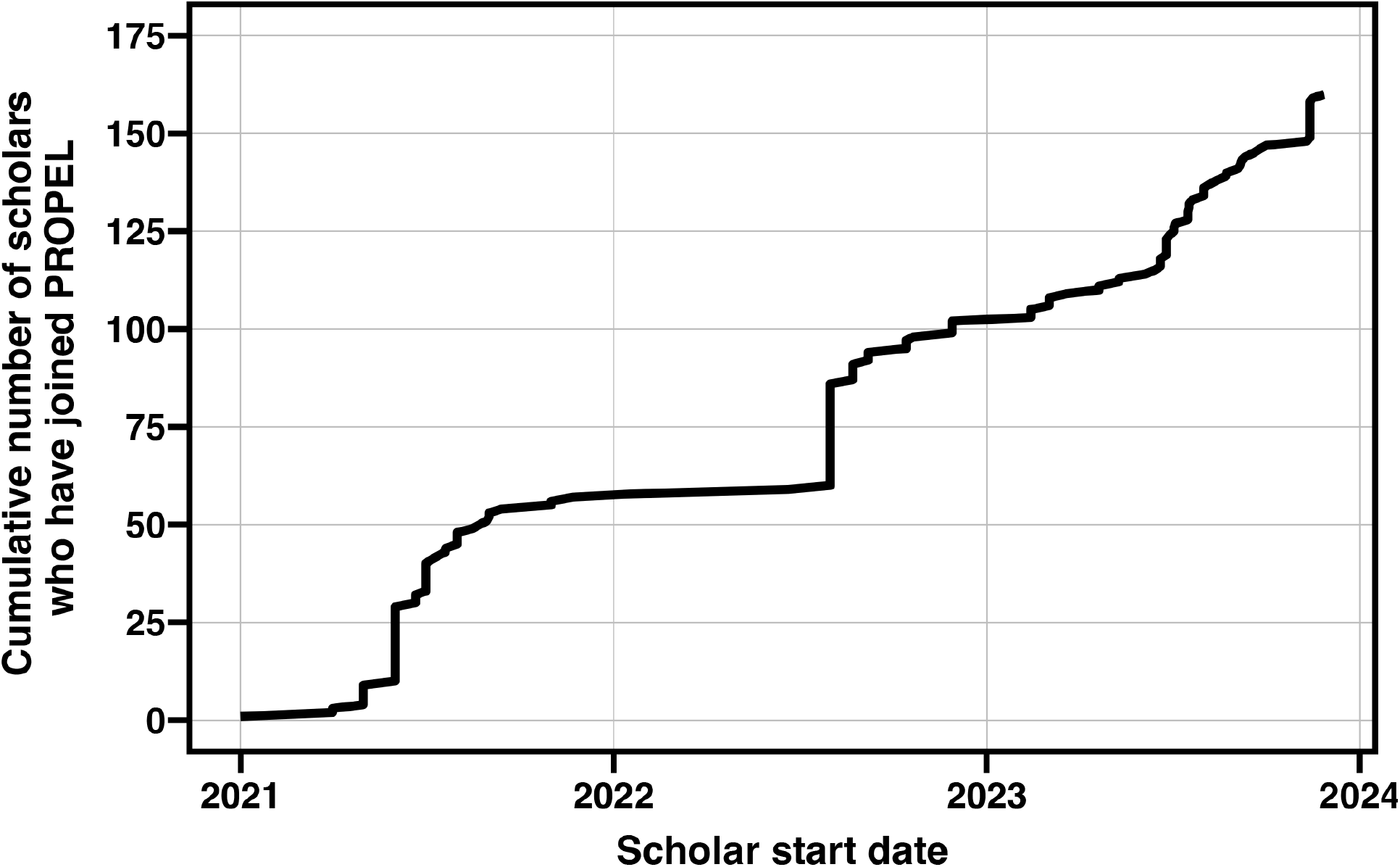
Graph showing the cumulative number of scholars who joined the PROPEL program from January, 2021 to December, 2023.

### Curriculum

The overall goal of the PROPEL curriculum is to provide scholars with scientific and professional development experiences that will help prepare them for success in a biomedical science PhD program. Since PROPEL scholars are also full-time employees, paid by their host lab, the curriculum is designed to complement their duties and hours in the lab. Specifically, curriculum events typically occur about three times per month and last for 1-2 hours per event. PROPEL mentors are aware of this commitment and, as part of the application process, mentors confirm that their scholar will be able to take time during the work week to participate in the curriculum fully. The PROPEL curriculum supports the growth of the scholar in four broad categories: (1) science communication; (2) critical thinking and rigorous application of the scientific method; (3) professional development; and (4) graduate program insight and preparedness. Below, the main components of the curriculum are described in more detail.

#### Literature Review

The Literature Review course guides the scholars through the process of reading, understanding, and discussing primary literature using a combination of lectures, assigned readings, and scholar presentations (Supplemental Material 1). It was initially designed to be taken by both first and second-year scholars but is now taken by only first-year scholars. Scholars meet for one lecture and one discussion section each month for ten months (September to June), and each month focuses on a different field of biomedical research. The faculty course director selects a paper for discussion each month, and a UCSF postdoctoral fellow with expertise in the area is recruited to give a lecture that provides the scholars with an introduction to the selected topic. A pair of scholars who are assigned to present the chosen article work closely with a coach (usually a UCSF PhD student or postdoctoral fellow) to take a deep dive into the article and prepare a “journal club” style presentation. They give the presentation at the discussion sessions, which serve as catalysts for peer engagement and critical analysis, guided by the facilitating coach, who ensures a robust exchange of ideas and offers constructive feedback to the presenters.

#### Data Science

The Data Science course uses lectures and hands-on group work to provide scholars with instruction in study design, setting sample sizes, choosing and performing statistical tests of hypotheses, and reproducible computational research (Supplemental Material 2). The course also provides a basic overview of R programming skills for data management, visualization, and analytics. These skills provide a foundational understanding of concepts that a majority of scholars will use in their graduate careers. The course meets once a week for eight weeks and is taken during the first year of the program.

#### Build Your Research Community

Build Your Research Community is a new course that we added at the beginning of the 2023-24 academic year. It is a free online course offered by iBiology (18, 19). It is designed to guide scholars through the steps of identifying mentors, particularly primary research mentors, and building and maintaining mentoring relationships. Scientists from a variety of backgrounds give concrete steps and strategies on how to build a mentoring network. In this course, participants create a detailed plan to choose a research group, dissertation committee, and build a mentoring network for greater success in graduate school; and learn techniques and strategies for establishing and maintaining healthy and professional relationships during graduate school, assessing their preferred communication and mentoring styles, and aligning expectations and goals for their research training. The course consists of six asynchronous online learning modules and five synchronous, in-person discussion sections and is taken during the second year in the program.

#### Graduate Application Preparation Series

This series of workshops supports scholars as they prepare their applications for graduate school (Supplemental Material 3). The workshops are open to all scholars and are not a required part of the curriculum but are primarily attended by scholars who plan to submit their applications in the current application cycle. The series includes a Graduate Admissions Panel, which is a facilitated conversation with graduate program directors or admissions committee experts who discuss what they are looking for when selecting successful applications and answer questions from scholars about the process; a Personal Statement Workshop, which provides scholars with one on one support for writing and revising their personal statements for the graduate application; and Mock Interviews, which gives scholars who have been invited for interviews with one or more graduate programs the opportunity to practice interviewing with UCSF faculty who are knowledgeable about the admissions process.

#### Secondary Mentors

All scholars have a primary mentor who hired them and serves as their direct supervisor. Additionally, each scholar is paired with another UCSF faculty member who acts as their secondary mentor (Supplemental Material 4). The goal is to offer scholars an opportunity to expand their professional network by establishing a relationship with another faculty member who can provide scientific and career guidance. While many secondary mentors also serve as the primary mentor for a PROPEL scholar within their own research group, some do not. In such cases, the secondary mentor program serves the additional purpose of broadening the community of participating faculty. Scholars are required to meet with their secondary mentors at least twice a year to discuss their research progress, review strides made toward their career objectives, and address any questions or concerns they may have. The secondary mentors also play a role in ensuring that scholars meet the expectations of the program and assessing their readiness to attend graduate school. After each meeting, the scholar provides a summary using a standardized meeting report form, which is then submitted to the program directors for review. In the first implementation of this component of the curriculum, scholars were paired with their Secondary Mentor by the program, based on broad similarities in research interest (e.g. both the scholar and the faculty member were in the Cancer Biology field). This was inefficient and resulted in many matches that were unproductive. Thus, we modified the process and now ask the scholar to work with their primary mentor to identify 1-3 faculty members they would like to be paired with. Scholars may take many different factors into account when choosing a secondary mentor, such as a shared racial, ethnic, or gender identity, similar career trajectory, or shared scientific interests. Once the scholar has made their selections, the PROPEL office reaches out to the faculty on behalf of the scholar to arrange the match.

#### Curriculum evaluation

The program curriculum is dynamic and continuously evolving to meet the needs of the scholars. These needs are determined by conducting course evaluations after each course and gathering feedback from the PROPEL Scholars Council.

## Demographics

Since the enrollment criteria during the first three years of the program included a requirement that the scholar meet the suggested eligibility criteria for an NIH Research Supplement to Promote Diversity in Health-Related Research (whether or not they are actually funded by this mechanism), all scholars in the program during that time identified as either belonging to a racial or ethnic group that is underrepresented in science or coming from a disadvantaged background (as defined by the NIH), or both. We collected this and other demographic information on the program application and present the aggregated statistics on **Table 1**. These data indicate that the population of PROPEL scholars is very diverse, with broad representation across many different demographic categories. Notably, 25% of PROPEL scholars indicated that they belong to two or more of the demographic categories (underrepresented race or ethnicity, economically disadvantaged, and living with a disability) that are used to establish eligibility for an NIH Diversity Supplement, emphasizing the intersectionality of these categories. Analysis of the scholars’ educational background (**Tables 2 and 3**) revealed that, while PROPEL scholars come from a wide variety of undergraduate institution types and geographic locations, the large majority (82.5%) come from schools within California, and most have attended undergraduate institutions that are categorized as having very high or high research activity (63.1% and 26.3%, respectively).

**Table 1:** Scholar demographics (n = 160)

| Demographics | Percent (n) |
| --- | --- |
| Race or ethnicity |  |
| American Indian/Alaskan Native/Native Hawaiian | 2.5 (4) |
| Black/African American | 19.4 (31) |
| Hispanic/Latine | 45.6 (73) |
| Pacific Islander/Filipino/Hmong/Vietnamese | 20.0 (32) |
| Two of the above race or ethnicity categories | 4.4 (7) |
| Economically disadvantaged | 38.1 (61) |
| Person with a disability | 3.8 (6) |
| Intersectionality |  |
| Two of the three demographic categories above | 23.1 (37) |
| All three of the above demographic categories | 1.9 (3) |
| Gender |  |
| Female/woman | 58.8 (94) |
| Male/man | 36.9 (59) |
| Genderqueer/Gender non-conforming/Gender non-binary | 4.4 (7) |

**Table 2:**
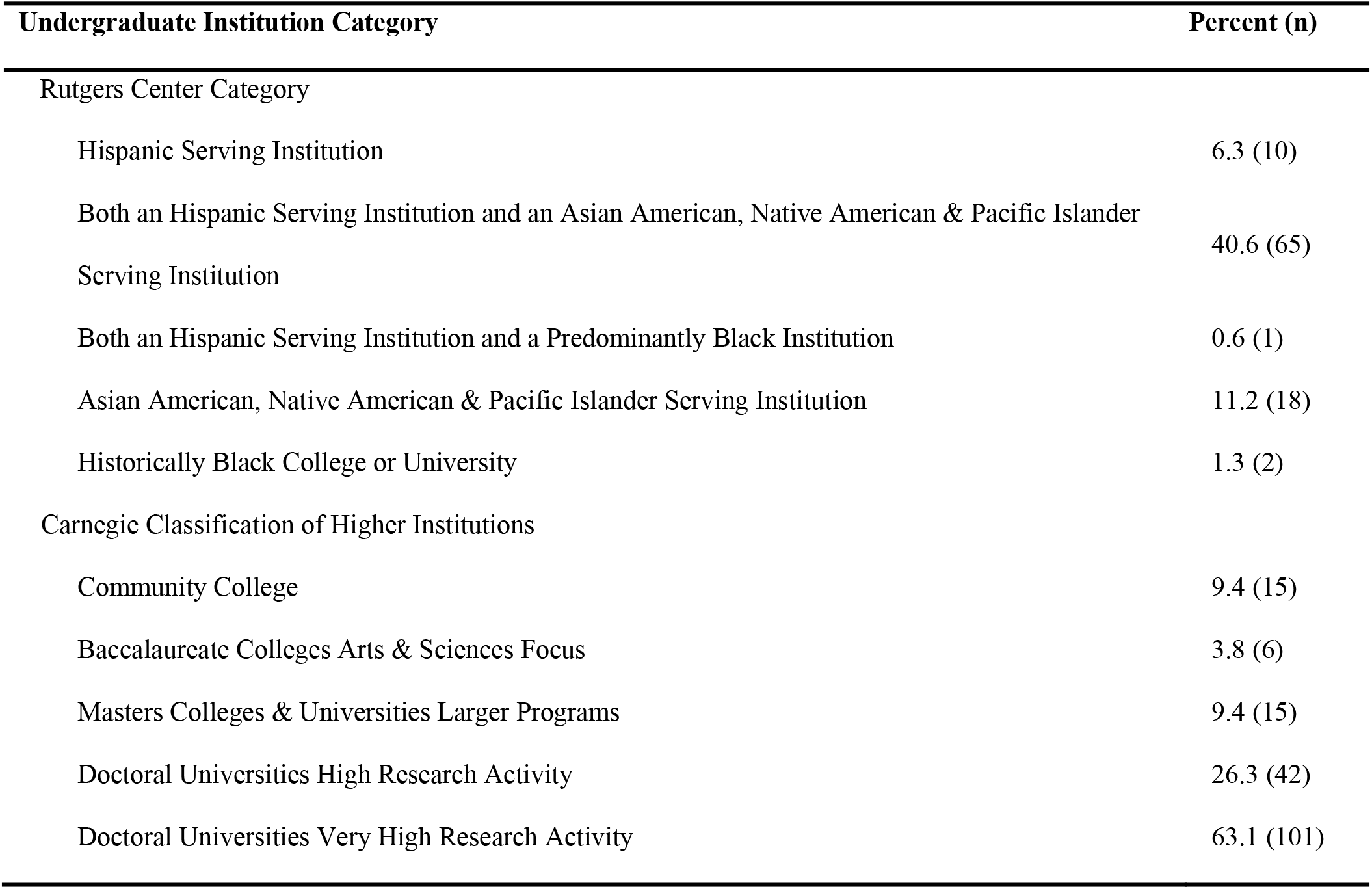
Scholar undergraduate institution type (n = 160)

| Undergraduate Institution Category | Percent (n) |
| --- | --- |
| Rutgers Center Category |  |
| Hispanic Serving Institution | 6.3 (10) |
| Both an Hispanic Serving Institution and an Asian American, Native American & Pacific Islander Serving Institution | 40.6 (65) |
| Both an Hispanic Serving Institution and a Predominantly Black Institution | 0.6 (1) |
| Asian American, Native American & Pacific Islander Serving Institution | 11.2 (18) |
| Historically Black College or University | 1.3 (2) |
| Carnegie Classification of Higher Institutions |  |
| Community College | 9.4 (15) |
| Baccalaureate Colleges Arts & Sciences Focus | 3.8 (6) |
| Masters Colleges & Universities Larger Programs | 9.4 (15) |
| Doctoral Universities High Research Activity | 26.3 (42) |
| Doctoral Universities Very High Research Activity | 63.1 (101) |

**Table 3:** Scholar undergraduate school location (n = 160)

| Undergraduate location | Percent (n) |
| --- | --- |
| California | 82.5 (132) |
| Other Western (WA, CO, and NV) | 3.1 (5) |
| Midwest and Southwestern (MO, MN, IL, and TX) | 5.0 (8) |
| Northeast and Mid-Atlantic (NY, PA, MA, RI, NY, PA, and DC) | 6.3 (10) |
| Southeast and the Caribbean (NC and PR) | 1.9 (3) |
| Mexico | 0.6 (1) |
| Online | 0.6 (1) |

A second enrollment criterion is that the scholar be employeed full time in the research group of a UCSF faculty member, but there were no restrictions based on the demographic or professional categories to which the faculty belonged. We present the aggregated statistics for faculty mentors in Tables 2-5. These data show that 13.2% of PROPEL mentors indicated that they identify as belonging to an historically excluded race or ethnicity, and an additional 8.8% were raised in an economically disadvantaged household or are a person with a disability (or both) (**Table 4**). By comparison, 15% of all UCSF faculty identify as belonging to an historically excluded race or ethnicity.^4^ The PROPEL program is also attracting faculty from all three career stages (Assistant, Associate, and Full Professor ranks) (**Table 5**) and a wide range of departments from all four schools (the Medical School, Dental School, Pharmacy School, and Nursing School) (**Supplementary Figure 1**). As a group, these faculty have substantial mentoring experience (**Tables 6 and 7**), as indicated by their prior experience across a range of career stages and membership in PhD programs.

**Table 4:** Mentor demographics (n = 114)

| Demographics | Percent (n) |
| --- | --- |
| Race or ethnicity |  |
| American Indian/Alaskan Native/Native Hawaiian | 0.9 (1) |
| Black/African American | 2.6 (3) |
| Hispanic/Latine | 7.9 (9) |
| Two or more of the above race or ethnicity categories | 1.8 (2) |
| Economically disadvantaged | 7.0 (8) |
| Person with a disability | 1.8 (2) |
| Intersectionality |  |
| All three of the above demographic categories | 0.9 (1) |
| Gender |  |

<sup>4</sup> https://diversity.ucsf.edu/sites/default/files/2023-12/ODO_Annual_Report_2022_2023_web.pdf

|  |  |
| --- | --- |
| Female/woman | 38.6 (44) |
| Male/man | 61.4 (70) |

**Table 5:** Faculty rank (n = 123)

| Rank | Percent (n) |
| --- | --- |
| Assistant Professor | 22.8 (28) |
| Associate Professor | 29.3 (36) |
| Full Professor | 48.0 (59) |

**Table 6:** Faculty mentorship experience (n = 123)

| Mentee stage | Percent (n) |
| --- | --- |
| Undergraduate | 79.3 (92) |
| Postbaccalaureate | 80.1 (93) |
| PhD | 87.1 (101) |
| Postdoctoral | 93.1 (108) |
| Any stage | 100 (116) |

**Table 7:**
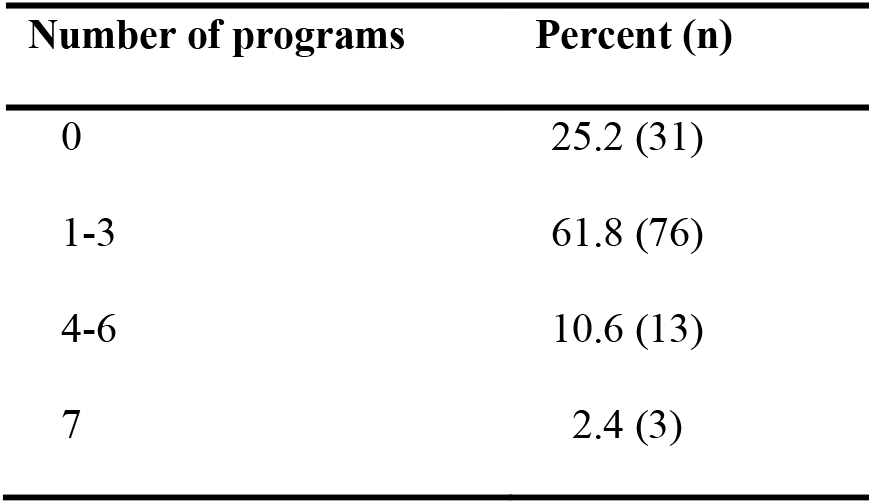
Faculty graduate program membership (n = 123)

| Number of programs | Percent (n) |
| --- | --- |
| 0 | 25.2 (31) |
| 1-3 | 61.8 (76) |
| 4-6 | 10.6 (13) |
| 7 | 2.4 (3) |

## Engagement

The PROPEL classes are not graded, and scholars are not awarded a certificate or degree for completing the program, so these incentives for participating in program activities do not apply. However, there is evidence that grading may not be effective either as an incentive for engagement or an accurate measure of motivation and ability, particularly among students from historically excluded backgrounds (20, 21). Instead, we impress upon the scholars when they join that these activities are provided for their benefit, and they are expected to participate as a condition of belonging to the program. We also communicate to the faculty the importance of allowing their scholars to attend PROPEL events at the scheduled times. In addition, we are careful to limit the number of required activities since we know that scholars have substantial obligations to their research group as a full-time, paid employee. We have found that this arrangement results in a high level of scholar engagement with the curriculum (**Table 8**).

**Table 8:**
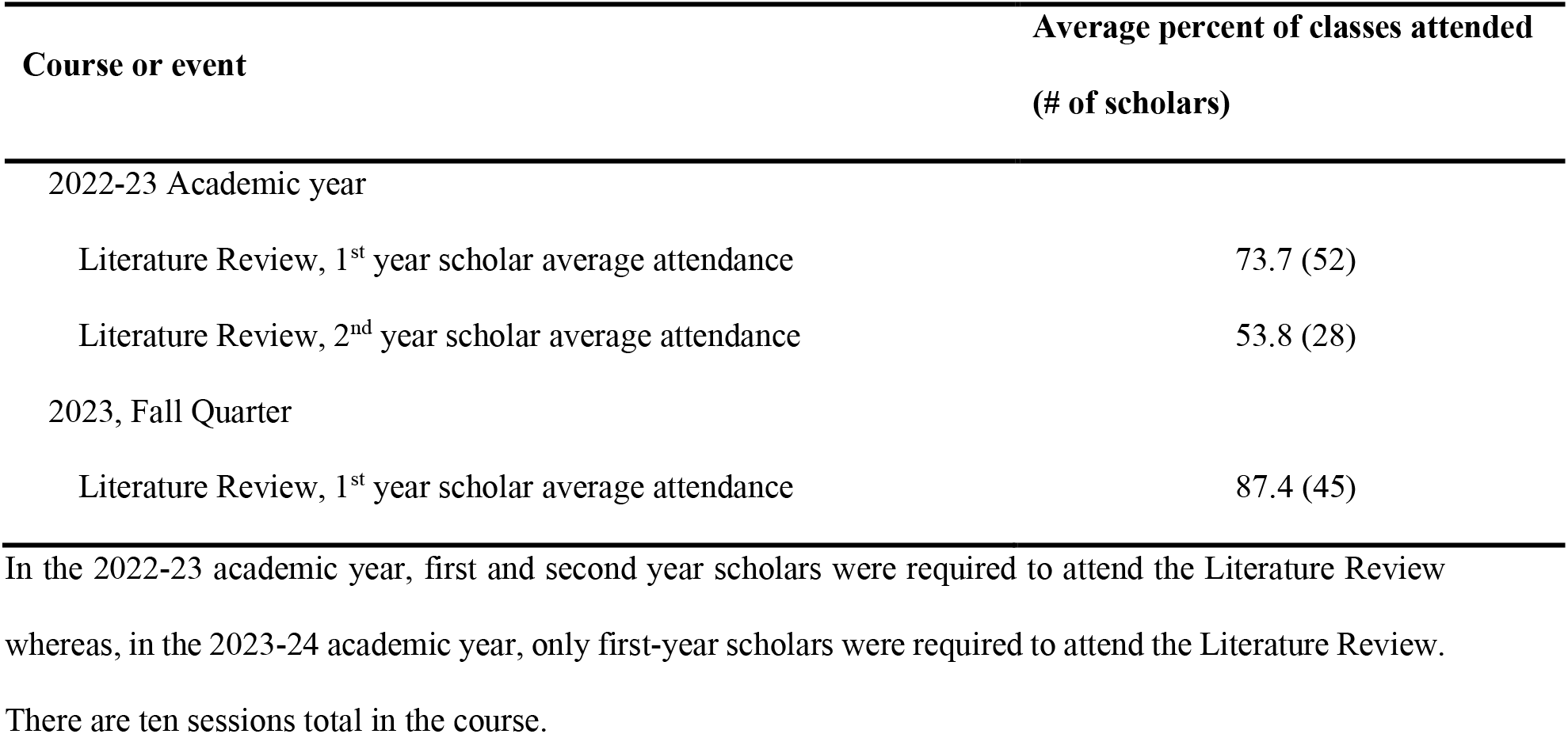
Average attendance per scholar at the Literature Review sessions.

In the 2022-23 academic year, first and second year scholars were required to attend the Literature Review whereas, in the 2023-24 academic year, only first-year scholars were required to attend the Literature Review. There are ten sessions total in the course.

## Outcomes

An enrollment criterion for the PROPEL program is that the scholar state an interest in pursuing a PhD or an MD/PhD degree. We include this criterion because we want to make it clear that the program is designed to train scholars for research-intensive careers, rather than, for example, an MD, DDS, or other professional degree that is not research-focused. Thus, in theory, all scholars have an interest in pursuing a research-intensive career when they enter the program. However, this is a formative period, and we recognize that some scholars may change their minds while they are in the program. In addition, scholars may elect to pursue opportunities at biotechnology companies that do not require a postgraduate degree. We present the aggregated statistics for the career choices made by all the scholars who left the program in 2022 and 2023 in **Table 9**.

**Table 9:** Attendance at other PROPEL classes and events.

| Course or event | Percent (n/total) |
| --- | --- |
| 2022-23 Academic year |  |
| Data Science, completed course and final project | 61.5 (32/52) |
| Graduate Admissions Panel, attended event<sup>2</sup> | 67.5 (54/80) |
| Annual Symposium, attended event |  |
| Faculty mentors | 35.6 (26/73) |
| Scholars | 90.8 (79/87) |
| 2023, Fall Quarter |  |
| iBiology BYRC, attended all 6 modules<sup>1</sup> | 48.0 (25/52) |
| iBiology BYRC, attended 2-5 modules<sup>1</sup> | 40.4 (21/52) |
| Graduate Admissions Panel, attended event<sup>2</sup> | 36.2 (34/94) |
| Personal Statement Workshop<sup>2</sup> | 33.0 (31/94) |

<sup>1</sup>In 2023, the iBiology course was only offered to the second-year scholars (52 in total).
<sup>2</sup>These events are mainly attended by second-year scholars, though first-year scholars are welcome to attend if they are interested.

**Table 9:**
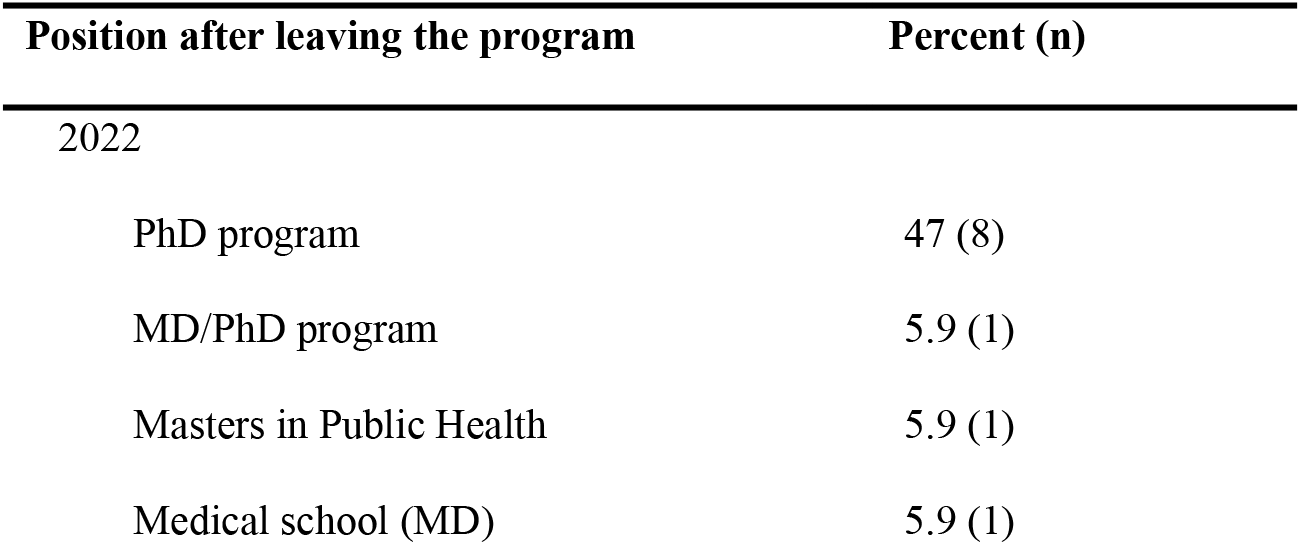
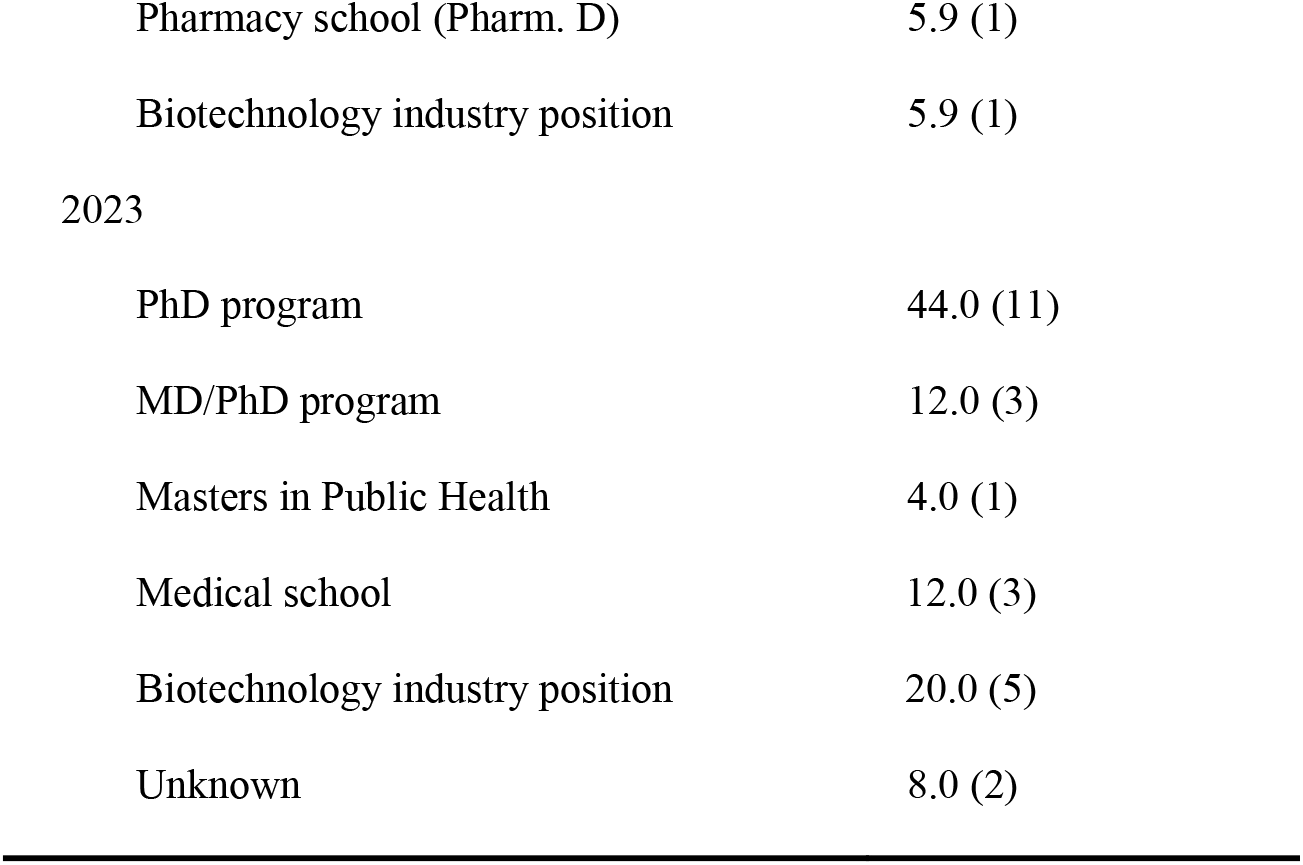
Scholar outcomes.

## Diversity supplement funding

Until 2023, one of the enrollment criteria for the PROPEL program was self-reported eligibility for support through the NIH Research Supplement to Promote Diversity in Health-Related Research, though funding through this mechanism is not required for enrollment. To support UCSF faculty who are interested in applying to this funding opportunity, the UCSF Research Development Office has compiled a website with useful information,^5^ including a password-protected library of successful applications, and the Office of Diversity and Outreach has developed a formalized process for confirming candidate eligibility.^6^ In addition, if the scholar’s application to the PROPEL program has been approved, the PROPEL program can provide a letter of support for the Research Supplement application.

## Institutional funding

Although the scholars are hired by their host lab, successful implementation of the program activities requires administrative support. In the first year of the program, this was provided by volunteer efforts from faculty and staff. However, as the program grew, it became important to have a dedicated program administrator who could coordinate all the logistics of the program, including admissions, curriculum, and scholar tracking. We were fortunate to be able to obtain generous financial support in the form of pilot grants from the UCSF Executive Vice Chancellor’s Office, the School of Medicine Dean’s Office, and local philanthropic organizations. In our experience, though there is some variation depending on the program size and the amount of time the program has existed, it requires approximately $2,000 per scholar per year to sustain the program operating budget.

## DISCUSSION

In the past three and a half years, the PROPEL program has scaled up to enroll over 150 scholars in total, with over 100 currently enrolled. This large pool of scholars has helped to build a critical mass on campus that makes the programming more vibrant and impactful, increases the visibility of the program, and builds community. The unique funding structure of PROPEL, in which scholars’ salaries and benefits are funded by their home lab, made it possible to support a large cohort on a relatively small programmatic budget. In addition, it allowed the program size to scale according to the resource availability in university research laboratories rather than a fixed number that is typical of programs supported by training grants. At its current scale, PROPEL has become, to our knowledge, the largest biomedical research program for postbaccalaureate scholars in the country. The fact that PROPEL has been able to grow so rapidly is a strong indicator of the demand for this type of programming and underscores the potential that the PROPEL model has to make a substantial impact toward reducing the inequities of the current system.

The primary goal of the PROPEL program is to prepare scholars for applying to graduate school and, indeed, over half of the scholars who left the program in 2022 and 2023 accepted an offer to a PhD or an MD/PhD program. However, one of the most important reasons for scholars to gain research experience before applying to graduate school is so that they have the first-hand experience to make an informed choice about whether graduate school is the right path for them. Thus, as expected, some scholars chose another path, such as attending medical school or accepting a position at a biotechnology company. We consider these to be successful outcomes as well. Future assessments of scholar outcomes will investigate the types of options scholars have when they leave the program and the motivations for choosing their next step.

When assessing the value added by a program like PROPEL, it is important to consider the role of biomedical research-focused postbaccalaureate programs within the broader context of all the different types of opportunities that help prepare scholars for graduate school as well as how scholars choose to join a postbaccalaureate program. Substantive research experience is essential for a competitive application to biomedical PhD programs (9) so, for students who did not obtain this type of experience as an undergraduate, enrollment in a postbaccalaureate program may be an excellent option. Interestingly, our finding that the large majority of PROPEL scholars completed at least a portion of their undergraduate studies at institutions with high or very high research activity (Table 2) suggests that, at least in some cases, the existence of these opportunities at an undergraduate institution is not sufficient to provide the necessary preparation for graduate school. Notably, even for scholars who do have a substantive research experience, working for one or more years as a full-time, paid professional researcher before applying to graduate school can be desirable for a variety of personal and professional reasons. Thus, postbaccalaureate programs such as PROPEL can help to fill this important need.

However, the existence of postbaccalaureate programs (as well as Masters programs) that are designed to help prepare scholars for PhD programs creates a risk that this type of experience will become an expectation of PhD program admissions committees, thus prolonging what is already a lengthy educational path to the PhD degree. With these concerns in mind, we emphasize in our outreach efforts and advertising materials that those scholars who feel ready should apply directly to PhD programs and view PROPEL only as a “backup” option. It is also important to note that scholars are not offered admission to PROPEL until after they are hired at UCSF, and only approximately half of the scholars in PROPEL participated in the matchmaking event. Thus, the majority of PROPEL scholars elected to accept a research technician position at UCSF through other channels for reasons that may or may not relate to the existence of the PROPEL program. Lastly, it is important to understand the efficacy of the PROPEL programming, and we are currently developing and administering validated surveys to assess this. However, the programming offered by PROPEL is much like the programming in a typical PREP program, which has been shown to be effective (15), so it is likely that the PROPEL programming is adding value in a similar manner. We also hope to investigate in future studies how much research experience PROPEL scholars had the opportunity to participate in as an undergraduate and why they chose to pursue their current position.

During the first three years of the program, one of the criteria for joining PROPEL was that the scholar self-identify as meeting the suggested eligibility criteria for an NIH Diversity Supplement. However, scholars can meet these eligibility criteria in different ways, and other demographic categories, such as gender and educational background, are not considered in this approach. Our analysis provides a baseline assessment of the PROPEL scholar demographics and reveals a breadth of diversity across all these categories. In addition, we found that PROPEL scholars have taken a range of educational paths, as indicated by the different types of universities they attended (**Tables 2 and 3**). In 2023, we removed the requirement that scholars must be eligible for a Diversity Supplement. We now consider all applicants who are US citizens or permanent residents and who meet the remaining described criteria, with priority given to applicants who clearly articulate how their personal history, achievements, and future career goals provide evidence of a strong commitment to promoting diversity, equity, and inclusion in science. Future assessments will reveal whether and how this change in admissions policy affects the program demographics.

In the past year, we have started to receive inquiries from faculty at other universities who are interested in starting a PROPEL program at their home institution. To help facilitate these efforts, we have created informational videos and documents that describe how to get started. In addition, we have compiled the educational materials we use in the PROPEL curriculum into a format that can be easily adapted for use elsewhere. These resources are available online^7^. We hope that, as new PROPEL programs are developed at other universities around the country, a national network of interconnected PROPEL programs will emerge, and this innovative model will become a new standard for postbaccalaureate education.

## Conclusion

The new model for a biomedical postbaccalaureate research program that we have developed with PROPEL in which scholars from historically excluded groups are recruited to UCSF through multiple means, including an online matchmaking event, hired by individual labs, and supported through university-sponsored programming is highly effective. At its current scale, the PROPEL program is well-positioned to significantly improve diversity in the biomedical research workforce. Future directions include consideration of expanding the eligibility criteria to include international scholars who would meet the other eligibility requirements and efforts to develop PROPEL programs at other universities.

---

**Supplemental Figure 1.**
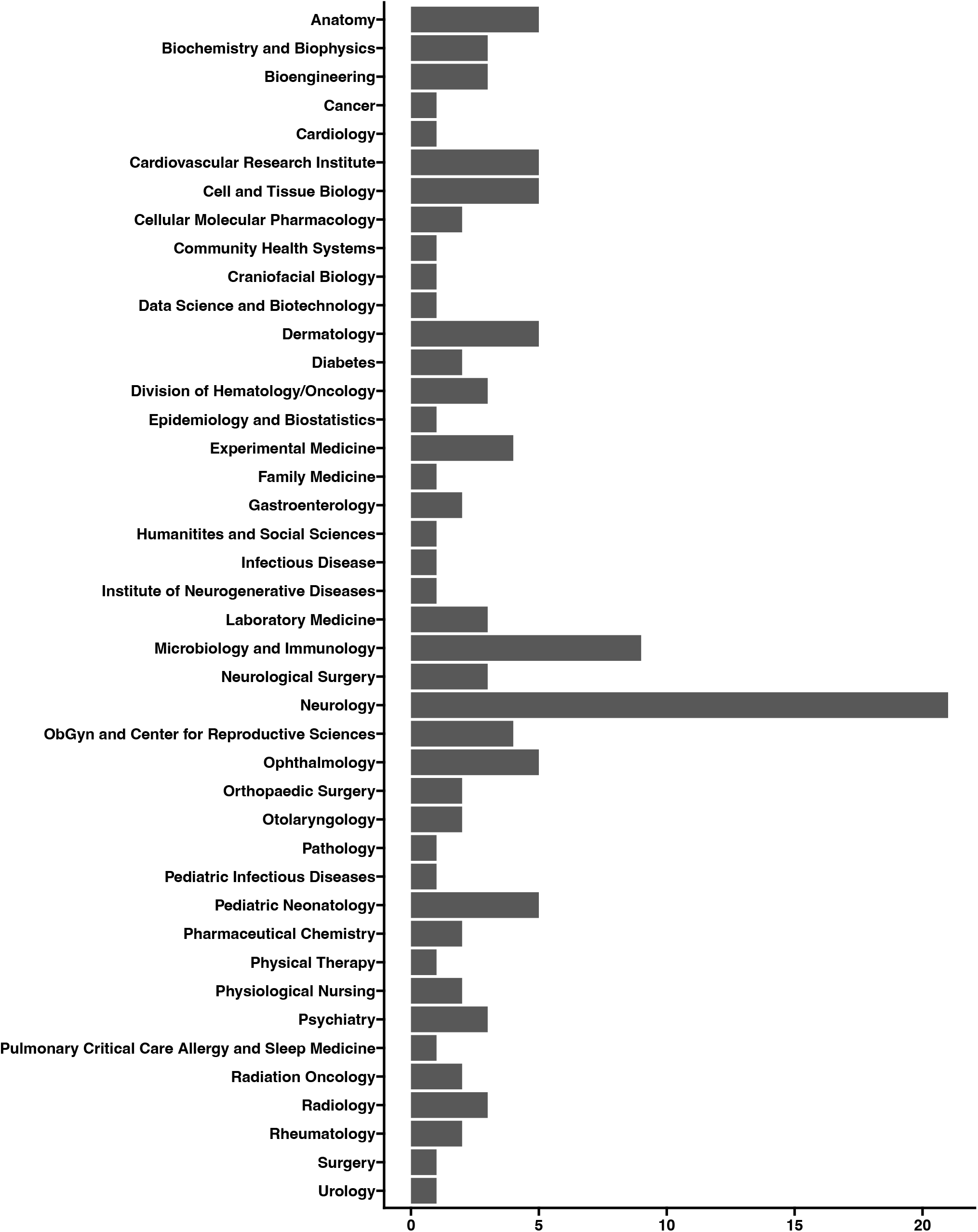

## Footnotes

1 https://graduate.ucsf.edu/admission/graduate-program-statistics/phd-program-statistics

2 https://www.ppic.org/publication/race-and-diversity-in-the-golden-state/

3 https://grants.nih.gov/grants/guide/pa-files/PA-23-189.html

4 https://diversity.ucsf.edu/sites/default/files/2023-12/ODO_Annual_Report_2022_2023_web.pdf

5 https://guides.ucsf.edu/rdo/diversitysupplements

6 https://diversity.ucsf.edu/programs-resources/grants-scholarships/diversity-supplements

7 https://propel.ucsf.edu/national-initiative

