## Supplemental Material 1 for "PROPEL: a scalable model for postbaccalaureate training to promote diversity in the biomedical workforce"

UNIVERSITY OF CALIFORNIA, SAN FRANCISCO

### Literature Review Course

PROPEL Program

##### Course Information

Faculty of Record: Claude Chapman

Professor

Office hours: [3-5pm Monday-Thursday]

Time:	Lecture: 4^th^ Thursday of the month preceding

Literature Review: Every 2nd and 3^rd^ Thursday of the month

Room:	Lecture: Zoom meeting [Link will be provided as needed]

Literature Review course: in-person Room# to TBA

##### Course Description

This course guides learners through the entire process of preparing a literature review and analyzing existing literature. Most importantly, the course develops skills in using evidence to create and present an engaging and critical analysis of science articles.

##### Course Objectives

The following course objectives describe what learners should be able to do by the end of the course.

- Critically assess evidence from published studies
- Increase exposure to rapidly evolving medical/science literature
- Be able to interpret published studies
- Develop communication skills
- Help inform in cutting edge methodologies

##### Course Communication

Please send your questions, requests, concerns to:

Claude Chapman, Director:

Jessica Allen, Program Administrator:

##### Teaching Method

This course utilizes a combination of lectures, assigned readings, and presentations by student.

Course Materials

Journal article readings will be provided by the 15^th^ of the preceeding month of the assignment and will be linked on the Propel website [https://propel.ucsf.edu/scholar-resources]

##### Course Schedule

| **Month** | **Topic** | **Key Activities** |
| --- | --- | --- |
| August | Introduction | - Class Zoom meeting: How to critically read and present a paper for journal club - Science Topics 101’s: Targeted Genetic Manipulation |
| September | Targeted Genetic Manipulation (e.g. CRISPR-Cas9, viral targeting, shRNAs) | - In Person: Journal club - Science Topics 101’s: In vivo disease modeling |
| October | In Vivo Disease modeling (e.g. transgenic systems, PDX | - In Person: Journal club - Science Topics 101’s: Molecular Mechanisms |
| November | Investigating Molecular Mechanisms | - In Person: Journal club - Science Topics 101’s: Transcriptomics |
| December | Break | No Journal Club  Happy Holidays! |
| January | Transcriptomics | - In Person: Journal club - Science Topics 101’s: Proteomics |
| February | Proteomics | - In Person: Journal club - Science Topics 101’s: Structural Biology |
| March | Structural Biology | - In Person: Journal club - Science Topics 101’s: Advanced microscopy and image analysis |
| April | Advanced Microscopy and Image Analysis | - In Person: Journal club - Science Topics 101’s: Artificial Intelligence in Science |
| May | Artificial Intelligence in Science | - In Person: Journal Club - Science Topics 101’s: Scientific Integrity |
| June | Scientific integrity | - In Person: Journal Club |

##### Graded Activities to complete Literature Review Course

| **Activity** | **Exception** |
| --- | --- |
| Participation to introduction session | Mandatory |
| Participation at 101’s lectures. | One excuse per year |
| Participation at Literature Review in person | Two excuses per year |
| Review from TA evaluating preparedness and contribution to Literature Review presentation |  |

##### 101’s sessions

These sessions are designed to provide scholars and Tas with background information on the selected topic relevant to assignment. They are lecture-based to facilitate the understanding of the selected article.

##### Assignment

Two first-year Propel scholars will be paired to present the selected article during the in-person Literature Review sessions. To the best of our ability, the students will be assigned to one science topic of her/his interest or the closest to their current research project.

##### Class Participation & Absence

Participation to Introduction lecture, 101’s lectures, and Literature Review sessions are mandatory. Participation will be recorded.

One excused participation for the 101’s lectures and two excused participations for Literature Review in-person sessions are allowed. A valid excuse needs to be granted by Dr. Chapman. Student needs to contact Dr. Chapman as soon as they know they will be missing a session and provide a reason. However, student can also request to participate in any other sessions that are ran in parallel.

##### Syllabus Revision

If the syllabus is revised, students will be notified by email. In addition a post will be made on Slack.

##### Mutual Respect and Humility

A key resource to student and faculty scholarship is the respectful exchange of ideas. At times issues or comments may surface that can cause offense or be hurtful to others. They may be made by either students or faculty, or sometimes raised in reading assignments. Such comments need to be discussed. It is important that we all commit to addressing issues as they arise. Please let faculty know if you are concerned about anything said in class or raised in the readings.

One process used in our School community to facilitate discussions is ***HEALS***:

**Halt** – Halt the discussion. Options include;

- Pause to consider the comment. Ask for clarification.
- Express appreciation for raising the issue.
- Use “I” language to express concern
- Focus on the idea. Deconstruct the comment without placing the individual on the defensive.

**Engage** with the issue

- Self check, check the room, look for body language.
- Go there. Discuss the issue.
- Let's talk about the issues embedded in this concept.
- What are the health care implications?

**Allow** exchange of opinions, stories, perspectives, and reactions.

- Let others express their thoughts, beliefs, feelings, and opinions.

**Learn** – Listen to one another through engaged and active listening

- Can we learn from one another’s experiences or observations?

**Synthesis** – Connect the dialogue to health equity/ quality of care

- How did the discussion itself work? Allow for opportunity to talk more later.

##### Student Resources

##### Technology & Library

- [Student IT Support](https://it.ucsf.edu/how-to/student-technology-support)
- [Ask a Librarian](https://www.library.ucsf.edu/contact)
- [Library Class Schedule](https://calendars.library.ucsf.edu/calendar/events)
- [Library Help Center](https://ucsflibrary.zendesk.com/hc/en-us)

##### Student Services

- [Student Portal](https://saa.ucsf.edu/studentportal/)
- [Student Disability Services](https://sds.ucsf.edu/)
- [Office for the Prevention of Harassment and Discrimination](https://ophd.ucsf.edu/)
- [Learning Resource Services](https://learn.ucsf.edu/)
- [Diversity and Outreach](https://diversity.ucsf.edu/)
- [Student Success Center](https://success.ucsf.edu/)
- [Student Health](https://studenthealth.ucsf.edu/)
- [Student Veteran and Military Support Services](https://veterans.ucsf.edu/)
- [International Students](https://isso.ucsf.edu/)
- [SON Student Editors](https://courses.ucsf.edu/course/view.php?id=1471)

##### UCSF Nondiscrimination Policy

University of California policy prohibits discrimination, including harassment, on the basis of race, color, national origin, religion, sex, gender, gender expression, gender identity, gender transition status, pregnancy, physical or mental disability, medical condition (cancer-related or genetic characteristics), genetic information (including family medical history), ancestry, marital status, age, sexual orientation, citizenship, or service in the uniformed services, including protected veterans.

University policy also prohibits retaliation against any individual or person seeking employment for bringing a complaint of discrimination or harassment pursuant to this policy.

For additional information regarding this policy or to file a complaint, please go to <https://ophd.ucsf.edu/>
