## Supplemental Material 2 for "PROPEL: a scalable model for postbaccalaureate training to promote diversity in the biomedical workforce"

PROPEL data science curriculum

**Overview**

Students will participate in data science training over the course of two months through weekly lectures and project-based assignments. Lectures will cover best practices for conducting reproducible research, ethics, applied statistics, computer programming, interpretation, and communication of results (see Curriculum below). Lectures will be instructor-led but will encourage peer-to-peer learning and discussion and will focus on practical applications to real world data.

Assignments will be designed to reinforce concepts covered in lectures, as well as contributing toward the goal of completing a project over the duration of the training series. The students will be given several population-based health or biological datasets to choose from for their projects. They will form small groups in which they must collaborate to analyze the data and answer a research question. At the end of the training series, they will present their findings to the class and instructors.

**Curriculum**

Session 1: Best practices for designing and conducting reproducible research

Session 2: Ethical considerations in biomedical research

Session 3: Hypothesis testing and association versus causation

Session 4: Data types, statistical distributions, measures of center and spread

Session 5: Statistical programming part 1: data exploration with univariate statistics and plots

Session 6: Statistical programming part 2: association testing and regression models

Session 7: Interpretation and presentation of results

Session 8: Project presentations

**Projects**

Students will get hands on experience working with a real dataset. The datasets we will use are still TBD, but may come from the [UC Irvine](https://archive-beta.ics.uci.edu/ml/datasets) repository.
