## Supplemental Material 3 for "PROPEL: a scalable model for postbaccalaureate training to promote diversity in the biomedical workforce"

PROPEL Application Workshop Series

Fall 2023/Winter 2024

Objective: To offer support and guidance to scholars who will be applying to graduate school in fall of 2023. This workshop series offers two tracks to support scholars who are etiher interested in PhD or MD/PhD program. Supports include application panels (consisting of faculty that reside on graduate admissions committees), personal statement workshops, mock interviews, and other supports specific to MD/PhD applications (AMCAS application help). Scholars are allowed to attend all workshop, if they chose. The Panel Workshops (MSTP Panel or Graduate Application Panel) is recommended for scholars who are still exploring whether a PhD or MD/PhD is right for them.

**PhD Track:**

Graduate Application Panel

Thursday, Sept 28^th^ 9 – 11am

Mission Bay, Mission Hall MH1401 & MH1402

Description: Faculty and program directors for UCSF graduate programs will attend to answer scholars’ questions about admission logistics, differences between graduate programs, tips for crafting a competitive application, and more.

Personal Statement Workshop

October 19^th^ 4 – 5:30pm

Parnassus, Campus Library, CL221 & CL222
October 20^th^ 4 – 5:30pm

Mission Bay, Mission Hall, MH1400

Description: Scholars learn general overview of the types of essays required for a PhD application and best practices for crafting a compelling set of essays. After the presentation, scholars meet with their assigned graduate peer mentor to get personalized feedback on their essay drafts. *Scholars must complete a draft of their essay by Oct 9^th^ to give their peer mentor time to review!*

Mock Interviews

January – February 2024 (scheduled individually)

Virtual meetings unless otherwise scheduled in person

Description: Scholars that are offered an interview at a graduate program can sign up to perform a mock interview with a PROPEL faculty member. Every attempt will be made to pair scholars with faculty from a department that matches the grad program to which they applied.

**MD/PhD Track**

MSTP Panel

Thursday, Oct 5^th^ 9 – 11am

Mission Bay, Genentech Hall N114

Description: Program director, Aimee Kao, will be in attendance to answer questions about UCSF MSTP and MD/PhD program in general. Also in attendance will be program manager Tiffiani Quan and several current MSTP graduate students. Scholars are encouraged to ask questions about MSTP programs, learn how an MD/PhD is different, learn about the application cycle, strategies around essay craft, and more.

Demystifying the AMCAS Application

Monday, Sept 25^th^ 9 – 10:30am

Remote

Description: Learn about the AMCAS application in general, a general timeline to apply to MD/PhD programs and more. This session is hosted by the Medical School Postbac Program

MSTP Essay Writing Workshop

Monday, Oct 9^th^ 4 – 5:30pm

Remote

Description: Learn how to craft your essays to tell a compelling and consistent narrative, along with the function of each essay. Get tips and tricks on where to begin when drafting each of the essays required on your AMCAS application. This session is hosted by the Medical School Postbac Program

AMCAS Nuts and Bolts

Tuesday, Feb 20^th^ 11:30 – 1pm

Remote

Description: At this session, hosted by the Medical School Postbac Program, engage in a deep-dive into the AMCAS application, including suggested timeline, explanation of each section, best practices, and more.
