## Supplemental Material 4 for "PROPEL: a scalable model for postbaccalaureate training to promote diversity in the biomedical workforce"

UCSF PROPEL

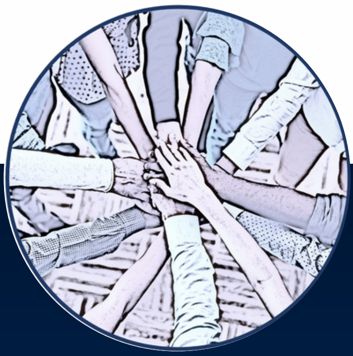

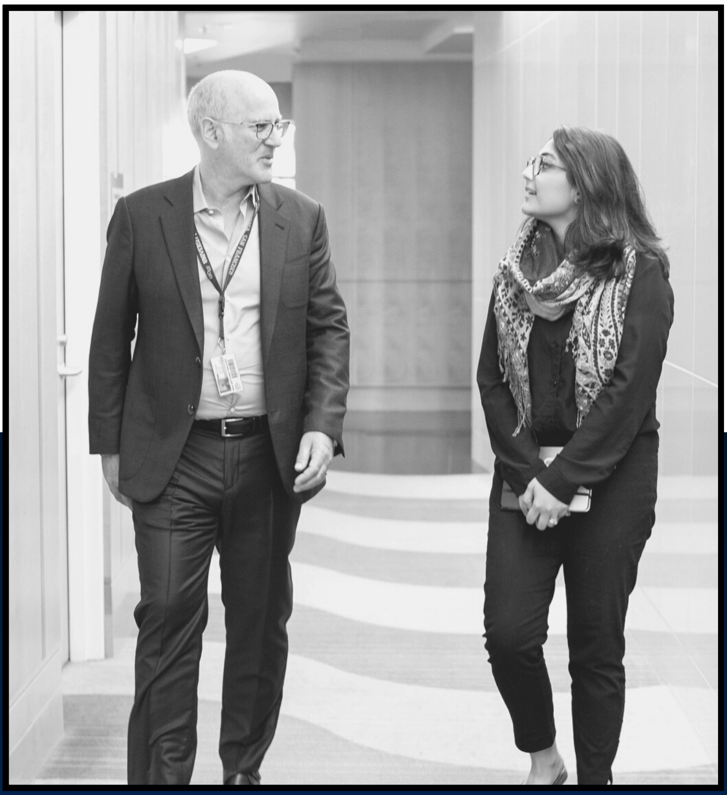

### MENTORSHIP GUIDEBOOK

GUIDEBOOK
GUIDEBOOK
GUIDEBOOK

FOR SCHOLARS & SECONDARY MENTORS

### TABLE OF CONTENTS


01
Mentorship: A Pillar of PROPEL

02
Benefits of a Secondary Mentor

03
Expectations

04
Where to Begin

05
Tips for Successful Mentorships

06
Troubleshooting

07
UCSF Resources

### MENTORSHIP: A PILLAR OF PROPEL

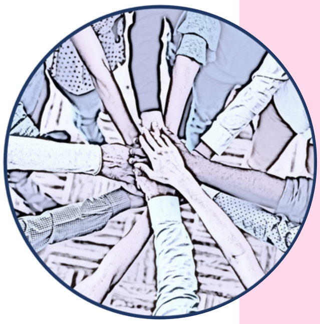

One of PROPEL's core tenents is strong mentorship. It is an important part of any career development, but especially at the post baccalaureate stage. Scholars at this stage are at a cross-roads; hoping to navigate career decisions while advancing in the field of science. Mentors serve as a guide to help scholar navigate these complicated and personal journeys, normalize struggles, identify and remove career roadblocks, and act as sources of inspiration.

THE ROLE OF SECONDARY MENTORS

All PROPEL scholars are already paired with a primary research mentor, but we ask scholars to find a secondary faculty mentor as well. While secondary mentors can be an additional source of research support, they have a more important role as an outside perspective, thought partner, and even a role-model. Secondary mentors can be in a related field, or have a career path that's aspirational for the PROPEL scholar, or come from a similar background or life experience.

Our secondary mentors serve as a vital resource for our scholars and ensure the success of the entire PROPEL program.

### BENEFITS OF A SECONDARY MENTOR

**HAVING A SECONDARY MENTOR IN SCIENCE CAN PROVIDE NUMEROUS BENEFITS.**

#### 01 Career Guidance

A secondary mentor may offer additional career guidance, sharing experiences and advice on navigating the scientific landscape. A secondary mentor is particularly valuable for early-career researchers. It may help you develop a well-rounded perspective on potential career paths.

#### 02 Expanded Network

Working with a secondary mentor can expose you to a broader professional network, including collaborators, colleagues, and experts in the field. This additional networking opportunities can contribute to your career development and open additional doors.

#### 03 Add'l Research Expertise

A second mentor may bring different expertise and perspectives broadening the range of skills and knowledge available to you. Their diverse expertise can offer insights from different scientific disciplines or methodological approaches.

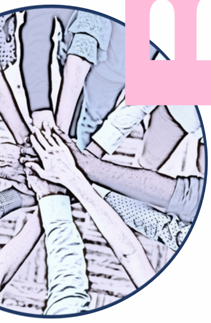

#### 04 Balanced Perspectives

Different mentors may have varying approaches to problem-solving and decision-making. A secondary mentor can offer another perspective, to help you make well-informed decisions.

#### 05 Interdisciplinary Collaboration

If the secondary mentor has expertise in a different field, it can foster interdisciplinary collaboration

#### 06 Work-Life Balance

The secondary mentor can offer guidance on work-life balance, stress management, and personal development. This type of support is essential for maintaining your well-being and overall satisfaction in their scientific journey.

### EXPECTATIONS

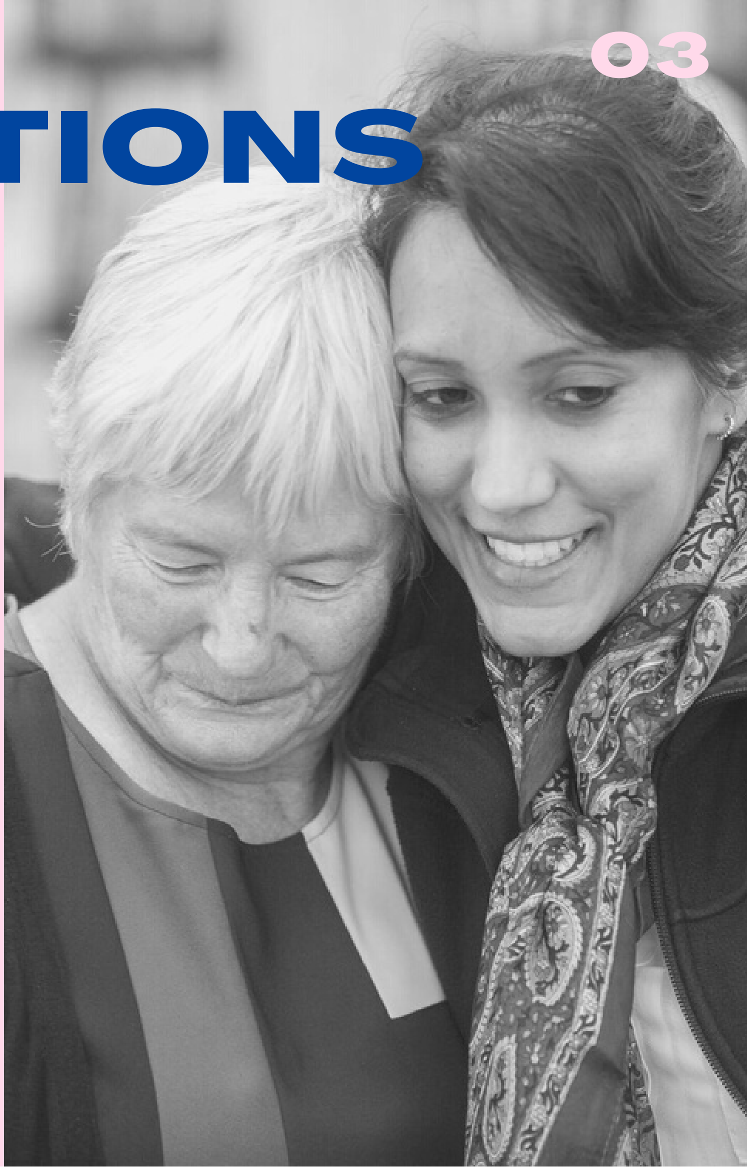

#### SECONDARY MENTOR EXPECTATIONS

- Provide active listening, guidance, and encouragement to the scholar.
- Connect the scholar with resources to help them achieve their goals.

#### SCHOLAR EXPECTATIONS

- Schedule meetings with the secondary mentor at least twice a year.
- Be open to receiving feedback and sharing your goals and challenges
- Prepare for the meeting using the Secondary Mentor Pre-Meeting Survey (below).

##### NUMBER OF MEETINGS 01.

PROPEL scholars are required to meet with their secondary mentor *twice per year*. Additional meetings are encouraged, at the discretion of both the scholar and secondary mentor.

##### DELIVERABLES 02.

Scholars are required to fill out a Secondary Mentor Pre-Meeting Survey before each meeting. A copy will be emailed to both mentor and scholar. In the survey, scholars record their research project, grad application progress, and career goals. After the meeting, scholars complete a Post-Meeting Survey, reflecting on the meeting.

##### RESOURCES 03.

As young scientists, PROPEL scholars are new to the scientific professional space and need resources to navigate this part of their career.

Secondary mentors are encouraged to act as a resource by sharing lessons learned on their own career journey, connecting scholars to colleagues and campus resources (page 6), as necessary.

Secondary mentors are also a resource for the PROPEL program, as you'll have insight into the scholars career development. If at any time you need more help, please don't hesitate to reach out to our program administrator at:.

### WHERE TO BEGIN

#### SETTING UP THE FIRST MEETING

Finding a time to meet can be challenging. Start by deciding what kind of meeting is most appropriate (zoom or in-person).

Offer several meeting options via email that work for you. Tip: checking Outlook's Scheduling Assistant is a great way to check someone else's calendar to find possible meeting times that match. Or create a doodle poll 1-on-1 with suggested times to find a match.

#### WHAT TO EXPECT AT YOUR FIRST MEETING

The first meeting between secondary mentor and scholar is all about getting to know one another.  Mentors and Scholars: introduce yourself, a bit about your personal/professional life, why you decided to participate in PROPEL, etc.

At this meeting, be sure to discuss how often you'd like to meet (PROPEL requires 2 meetings per year).

#### WHAT TO ASK AT YOUR FIRST MEETING

Here some topics to discuss at your first meeting:

- For scholars: Clarify the role you'd like your mentor to have
- For mentors: clarify the role you can provide (eg: encourager, advisor, professional development coach, etc.)
- Go over the Secondary Mentor Pre-Meeting Survey (emailed before the meeting).
- Schedule your next meeting

#### INSTRUCTIONS FOR MENTORS

Remember that you are resource for the scholar, both for career development and for navigating their existing career in science. A great way to start? Asking questions like: "How can I help?"

### TIPS FOR SUCCESSFUL MENTORSHIPS

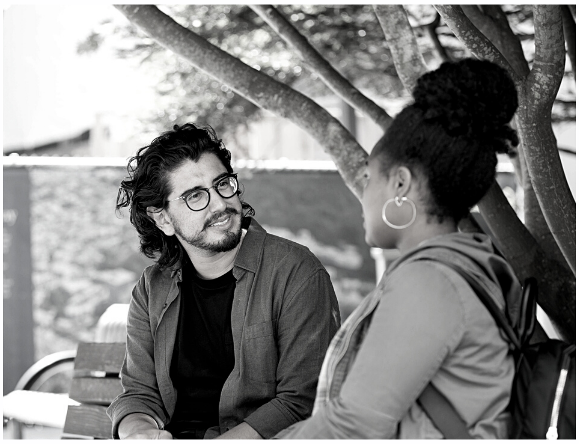

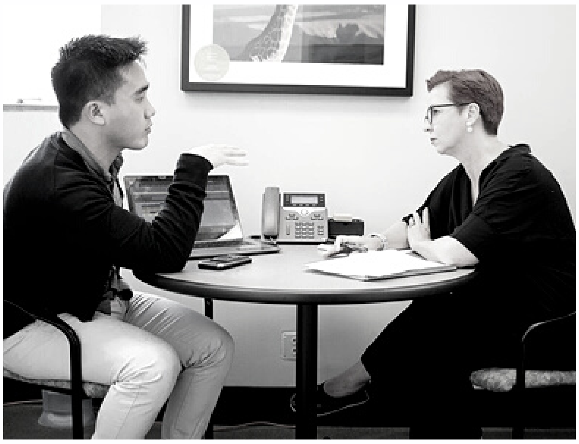

#### For Mentors

- Use active listening.
- Share your successes and failures. This will help your scholar feel comfortable sharing about their own experiences.
- Don't assume anything about your scholar or their life experiences.
- Be clear about your availability. Mentoring is volunteer work, and we want to be respectful of your donated time, so be clear about what you are able to do.

#### For Scholars

- Prepare for meeting with the Secondary Mentor Pre-Meeting Survey and be prepared your responses at the meeting.
- Be open to sharing challenges as well as successes. Secondary mentors can be a great place to get help navigating your professional career or guidance on difficult topics.
- Be clear about what you want from your mentor and what you find helpful.
- You don't always have to take your mentor's advice, but be sure to listen fully before making a decision.

### TROUBLESHOOTING

**Mentorships many not always work out and that's ok. Here's what to do if you're having trouble.**

##### Meetings difficult to schedule/email response slow

Make sure to check in about communication preferences. Sometimes a slack message or a phone call is preferred to email. Many faculty have assistants that manage their calendars. Be sure to discuss if this is the case.

We all get busy, so sending a follow-up meeting after one week with a short reminder can keep meeting requests from getting lost.

##### Shifting career paths/availability

.Scholars are at the very beginning of their career path. Therefore their career goals can and do shift, and that OK. Sometimes its a change in interest in a particular field of study, or about graduate school, or which degree to pursue (PhD vs MD/PhD). It can mean that what once was a good secondary mentor match no longer fits. If there is a shift in career goals, and it's OK to ask to rematch with someone that shares these new interests and aspirations.

Similarly, faculty commitments and time availability can change. If a secondary mentor's schedule becomes too tight to offer adequate support, it's OK to ask for a rematch to someone who has more time.

#### WHAT TO DO IF IT'S NOT WORKING...

Discussing early how to check-in on the status of your mentor/scholar relationship will help facilitate those conversations later if something isn't working. If the match doesn't seem quite right, we can help! Contact Jessica Allen at for help, or to find another match.

### UCSF Resources

**Student Success Center**
success.ucsf.edu

**Student Life**
studentlife.ucsf.edu

**Basic Needs**
basicneeds.ucsf.edu

**Student Diability Services**
sds.ucsf.edu

**Student Health & Counseling**
studenthealth.ucsf.edu

**Learning Resource Service**
learn.ucsf.edu

**Multicultural Resource Center**
mrc.ucsf.edu

**LGBT Center**
lgbt.ucsf.edu

**Student Financial Services**
finaaid.ucsf.edu

**Campus Life Services**
(Hoursing, Transportation, Fitness & Rec, Family Services)
campuslifeservices.ucsf.edu

**Undocumented Student Support Services**
undocu.ucsf.edu

**Office of Career & Professional Development**
career.ucsf.edu

**Student Veteran & Military Support Services**
veterans.ucsf.edu

**CARE Advocacy Resources & Education**
careadvocate.ucsf.edu
